# Prominent features of the amino acid mutation landscape in cancer

**DOI:** 10.1101/136002

**Authors:** Zachary A. Szpiech, Nicolas B. Strauli, Katharine A. White, Diego Garrido Ruiz, Matthew P. Jacobson, Diane L. Barber, Ryan D. Hernandez

## Abstract

Cancer can be viewed as a set of different diseases with distinctions based on tissue origin, driver mutations, and genetic signatures. Accordingly, each of these distinctions have been used to classify cancer subtypes and to reveal common features. Here, we present a different analysis of cancer based on amino acid mutation signatures. Non-negative Matrix Factorization and principal component analysis of 29 cancers revealed six amino acid mutation signatures, including four signatures that were dominated by either arginine to histidine (Arg>His) or glutamate to lysine (Glu>Lys) mutations. Sample-level analyses reveal that while some cancers are heterogeneous, others are largely dominated by one type of mutation. Using a non-overlapping set of samples from the COSMIC somatic mutation database, we validate five of six mutation signatures, including signatures with prominent arginine to histidine (Arg>His) or glutamate to lysine (Glu>Lys) mutations. This suggests that our classification of cancers based on amino acid mutation patterns may provide avenues of inquiry pertaining to specific protein mutations that may generate novel insights into cancer biology.

## INTRODUCTION

Cancers have been described as open, complex, and adaptive systems [1]. Reflecting this, cancer progression is determined in part by genetic diversification and clonal selection within complex tissue landscapes and with changing tumor properties and microenvironment features [2, 3]. Genetic sequencing of tumor samples has been critical in developing the evolutionary theory of cancer. While cancers traditionally have been—and continue to be—classified by tissue of origin, genetic sequencing has allowed for classification based on driver mutations [4] or nucleotide mutation signatures [5]. However, cancer cell adaptation is mediated by changes at the protein level that alter cell biology and enable cancer cell behaviors such as increased proliferation and cell survival. Existing cancer classifications by nucleotide mutation signatures lack a link between the underlying genetic landscape and effects on cancer cell phenotypes. Analysis of cancers by amino acid mutations could provide important connections between cancer evolution and adaptive biological phenotypes as well as provide insight into how specific classes of amino acid mutations may generally alter the function of the proteins in which they are found. There have been some studies to examine amino acid mutations across cancers [6–8], but these have relied on simple mutation counting methods.

Here we take a machine-learning approach to analyze amino acid mutations across 29 cancers in order to identify characteristic amino acid mutation signatures. Our analyses reveal that some cancer types have mutation signatures dominated by arginine to histidine (Arg>His) mutations, some have signatures dominated by glutamate to lysine (Glu>Lys), and others have more complex signatures that lack a single dominant amino acid mutation. These signatures were further validated in a non-overlapping set of samples from the COSMIC somatic mutation database. Importantly, this approach identifies not only which amino acid mutations are prevalent among cancers but also which amino acid mutations tend to occur together. For example, cancers with strong Arg>His signatures will also frequently have many Ala>Thr mutations but are unlikely to have many Glu>Lys mutations (despite all of these amino acid transitions resulting from a G>A nucleotide mutation).

## RESULTS

### Several cancers are enriched for R>H and E>K amino acid mutations

Multiple studies have interrogated nucleotide mutation biases by analyzing somatic variation across a wide range of cancers [4, 5]. However, in protein coding regions of the genome (i.e. the exome), it is essential to study patterns of amino acid variation to reveal information about potential functional effects at the protein level. We characterized the global properties of amino acid mutations encoded by somatic mutations across a range of cancers by analyzing a tumor-normal paired mutation database [5] consisting of 6,931 samples across 29 cancer types. We applied filtering to remove sequencing artifacts and restricted mutation data to nonsynonymous amino acid mutations (see Materials and Methods, S1 Table and S2 Table for details).

Using this amino acid mutation database, we performed an unbiased characterization of mutation signatures across cancer types using Non-negative Matrix Factorization (NMF), which has proven to be a useful tool for pattern discovery in cancer tissue mutation datasets [5] and other biological systems [9]. Applying NMF to the pooled mutation data reveals six mutation signatures at the amino acid level (S1G Fig), including two with strong Arg>His components and two with strong Glu>Lys components (Fig 1A, S1 Fig). Although the cancers are comprised of a mixture of the signatures identified, ten cancers (AML, colorectal, esophageal, low grade glioma, kidney chromophobe, medulloblastoma, pancreatic, prostate, stomach, and uterine) have majority contributions from Arg>His-prominent mutation signatures (R>H and A>T/R>H). We also identify four cancers (bladder, cervix, head and neck, and melanoma) that have majority contributions from Glu>Lys-prominent mutation signatures (E>K and E>K/E>Q). Additionally, there are two complex signatures not dominated by any particular amino acid mutation. Glioblastoma, kidney papillary, liver, and thyroid cancers have majority contribution from the Complex 1 signature, and lung adenocarcinoma, small cell lung, squamous cell lung, and neuroblastoma cancers all have majority contribution from the Complex 2 signature. Finally, seven cancers from a variety of tissues (ALL, breast, CLL, clear cell kidney, B-cell lymphoma, myeloma, and ovarian) have heterogeneous mutation signature contributions.

**Fig 1.**
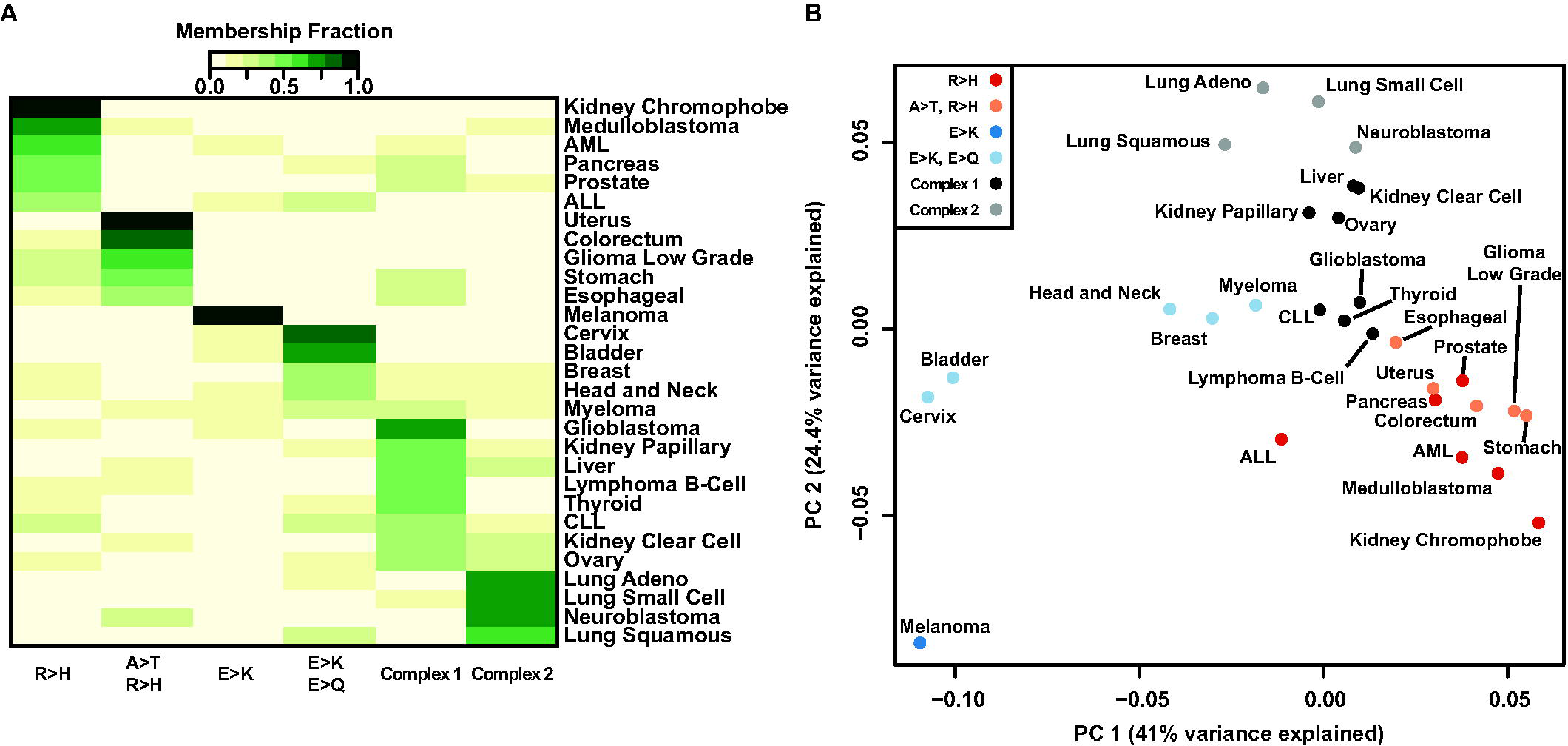
Arg>His and Glu>Lys mutations define mutation signatures of a subset of cancers. (A) Heatmap representation of six-component NMF clustering. Of the six amino acid mutation signatures identified, four have prominent charge-changing mutations: Arg>His (R>H), Glu>Lys (E>K), or Glu>Gln (E>Q). Two complex signatures were also identified. Color scale represents scaled contribution of each signature for a given cancer type. Signature and NMF fit details can be found in S1 Fig. (B) Principal component analysis of nonsynonymous amino acid mutations. PC1 separates cancers with high R>H from cancers with high E>K; PC2 separates cancers with complex signatures. Colors represent the greatest mutation signature contributing to a given cancer. Individual PC loadings can be found in S2 Fig.

### Visualizing Amino Acid Mutation Properties with Principal Component Analysis

To alternatively visualize the amino acid mutation spectrum, we use principal component analysis to reveal cancers clustering by dominant mutation classes (Fig 1B). We find that PC1 separates Arg>His dominant cancers from Glu>Lys dominant cancers and that PC2 separates cancers with more complex signatures (S2 Fig). This result reinforces our observation that Arg>His and Glu>Lys mutations are characteristic signatures of several cancers.

### Individual Cancer Samples Recapitulate Amino Acid Mutation Patterns

We also analyze samples individually with NMF and find that Arg>His and Glu>Lys features continue to dominate (Fig 2A and S3 Fig). For many cancer subtypes (melanoma, bladder, uterine, colorectal, low-grade glioma, cervix, neuroblastoma, and the three different lung cancers), individual patients within each cancer exhibit consistent amino acid signatures (Fig 2B). This is true even within clinically diverse cancers such as bladder, uterine, colorectal, and lung cancer, which all have multiple identified driver mutations. This suggests that the amino acid signatures we identified may be independent of underlying driver mutations and may instead be a consequence of common features of the cancer, tumor microenvironment, or selective pressures, all of which may be targeted therapeutically.

**Fig 2.**
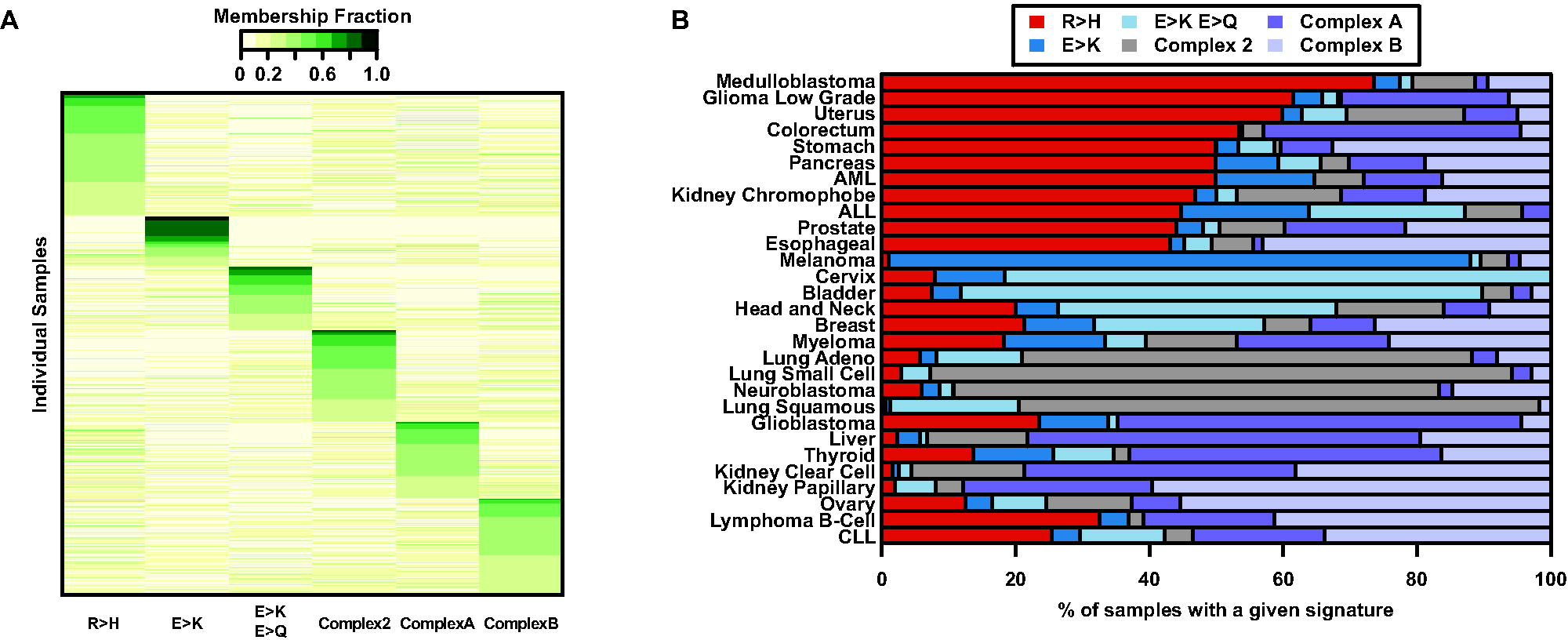
Amino acid mutation signatures for individual samples. (A) A heatmap representation of the six-component NMF clustering results for individual cancer samples (only those with >10 total nonsynonymous mutations). Samples with the same maximum signature component were grouped and sorted. Four amino acid mutation signatures identified (R>H, E>K, E>K/E>Q, Complex 2) overlap with signatures in Fig 1A. Color scale represents scaled contribution of each signature for a given sample. Signature and NMF fit details can be found in S3 Fig. (B) Bars show the total fraction of individual samples with a majority of a particular signature within each cancer. Within cancers, a large fraction of individual samples tend to have similar signature components.

As NMF decomposes a sample into a mixture of characteristic signatures, we can further visualize the normalized mixture coefficients from the individual-level NMF along the three mutation signatures with dominant Arg>His or Glu>Lys components (R>H, E>K, and E>K/E>Q signatures; Fig 3) to determine whether samples tend to be an equal mixture of several signatures or whether they tend to be exclusively composed of a single signature. Indeed, Fig 3 shows a clear separation of samples with a high proportion of Glu>Lys from other signatures.

**Fig 3.**
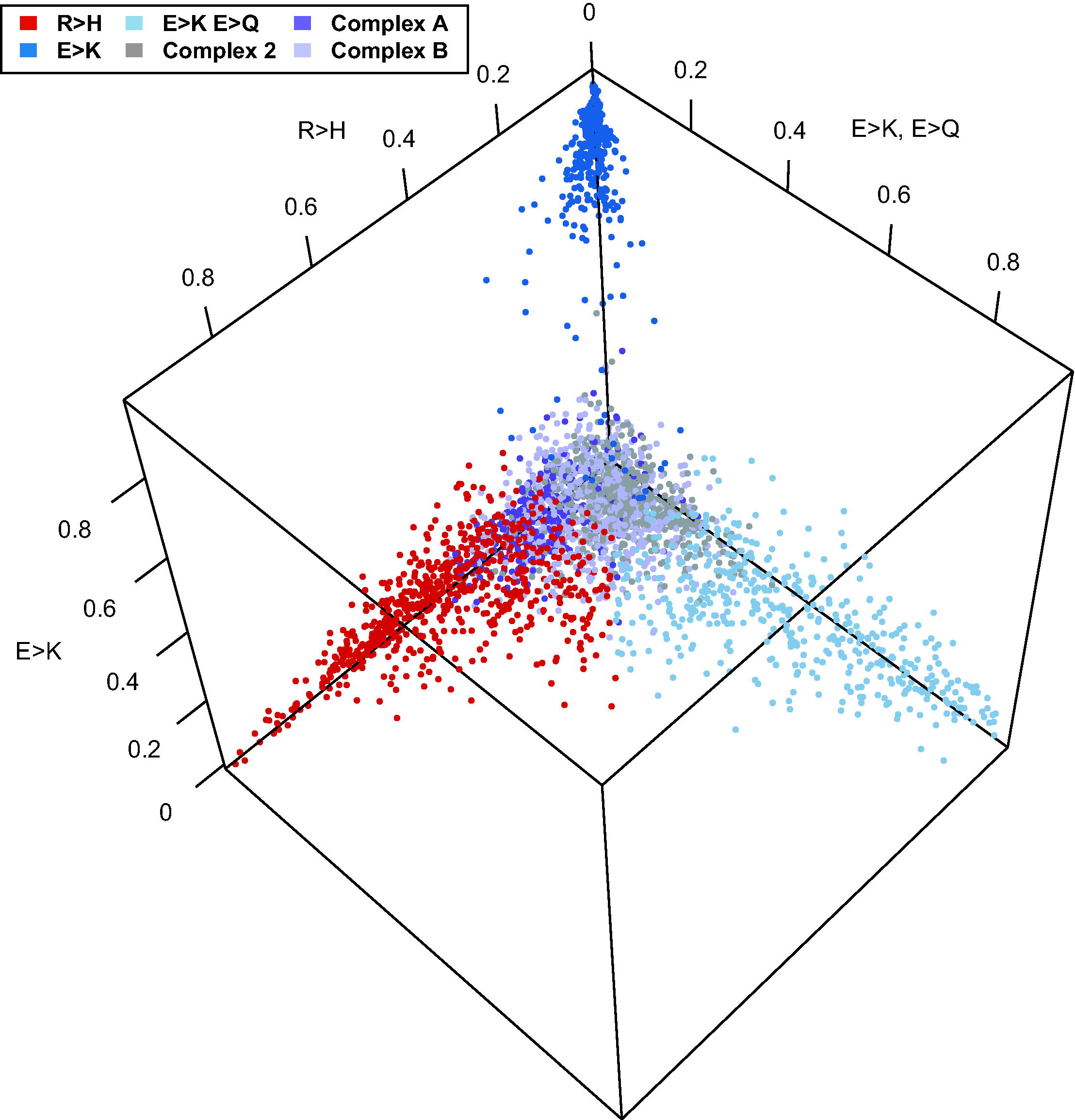
Normalized NMF mixture coefficients for individual samples. Plot of the normalized mixture coefficients across the three mutation signatures with high R>H or E>K components for every individual sample. Colors represent the greatest contributing mutation signature for each sample based on the full individual-level NMF analysis. Here we see a dramatic separation of samples in the E>K component to the near exclusion of other signatures.

### Mutation Signature Validation

We validated the NMF signatures with an orthogonal data set (see Materials and Methods) from the COSMIC database [10]. The six mutation signatures identified from COSMIC (S4 Fig) overlap substantially with previously identified signatures (S3 Fig). We calculated correlation coefficients between all COSMIC Data signatures and each Alexandrov Data signature. When the correlations are very high, this indicates that NMF has identified the same general mutation signature in the two different data sets. Indeed, we found high correlation between the COSMIC signatures and our initially identified signatures for five of the six (Fig 4): R>H, E>K, E>K/E>Q, Complex 2, and Complex A are replicated. The Complex B signature does not replicate as a separate signature, but appears to be largely incorporated into the other complex signatures. Interestingly, a new R>Q/R>W signature is identified as a separate component in the COSMIC data. On inspection of the Alexandrov R>H component we identified, we see that R>Q and R>W are prominent components. Increased sample size in our replicate data set likely enabled NMF to discriminate between these two signatures in COSMIC.

**Fig 4.**
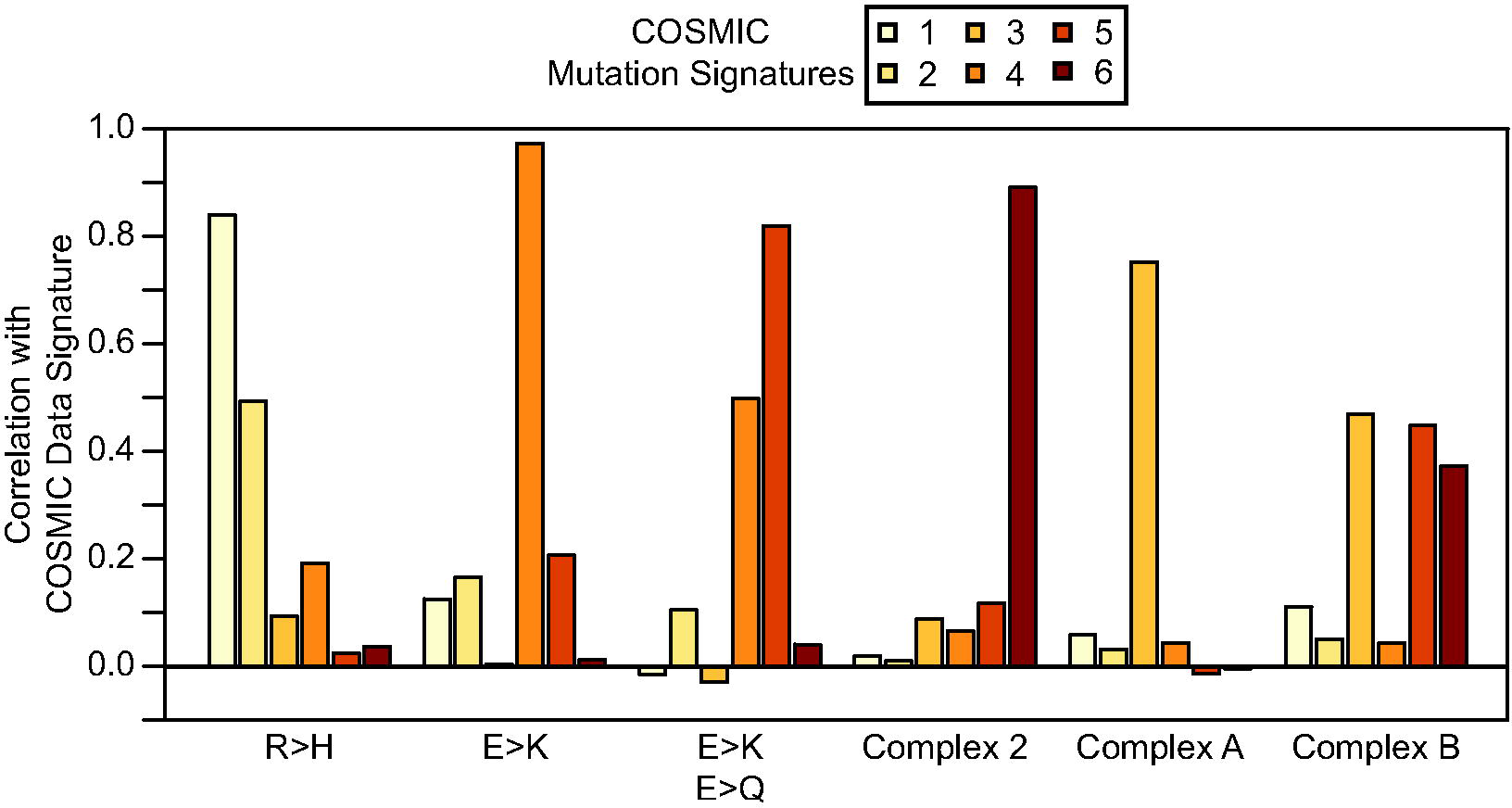
Correlations of COSMIC and Alexandrov mutation signatures. For each COSMIC mutation signature we calculated the correlation with each Alexandrov mutation signature. Five of six Alexandrov mutation signatures replicate in the COSMIC data for k = 6 mutation signatures. Alexandrov Signature R>H is replicated by COSMIC Signature 1, although a subset of mutation types that clustered in the original R>H are identified in the larger COSMIC data set as Signature 2. Alexandrov Signature E>K is replicated by COSMIC Signature 4. Alexandrov Signature E>K, E>Q is replicated by COSMIC Signature 5, although it is also correlated with COSMIC Signature 4 as each of these signatures share mutations. Alexandrov Signature Complex 2 is replicated by COSMIC Signature 6. Alexandrov Signature Complex A is replicated by COSMIC Signature 3. Alexandrov Signature Complex B is not faithfully replicated by any single COSMIC Signature, as the signal appears to be spread amongst COSMIC Signatures 3, 5 and 6.

## DISCUSSION

Proteomic changes can allow cancer cells to adapt to dynamic pressures including changes in matrix composition, oxygen and nutrient availability, intracellular metabolism, as well as increased intracellular pH (pHi), the latter enabling tumorigenic cell behaviors [11–15]. Our analyses reveal that a subset of all possible amino acid mutations dominate the mutation landscape of cancers, with Glu>Lys and Arg>His mutations being the most prominent features of identified mutation signatures.

Charge-changing mutations, whether buried or surface-exposed, can alter protein charge, electrostatics, and conformation [16]. Electrostatics of surface residues have been shown to play a key role in protein-protein interactions [17], protein-membrane interactions [18, 19], and kinase substrate recognition [20]. While it is important to note that our analyses are agnostic to the location of the mutation within the proteome and within a protein, the strong bias towards amino acid mutations that alter charge in our identified mutation signatures may suggest an adaptive advantage conferred by these mutations.

Glu>Lys mutations swap a negatively charged amino acid for a positively charged amino acid, which may in some cases affect protein function. Indeed, in some cases buried lysine mutations can have distinctly upshifted pKas [24] as well as induce global protein unfolding upon charging that alters mutant protein stability and function [21]. Furthermore, Glu> Lys mutations have been known to affect the function of PIK3CA [22–24].

Arg>His mutations swap a positively charged amino acid for a titratable amino acid. Whereas arginine (pKa ~12) should always be protonated, histidine (pKa ~6.5) can titrate within the narrow physiological pH range. Indeed, the pH-sensitive function of many wild-type proteins has been shown to be mediated by titratable histidine residues [25–27]. Moreover, recent work has shown that some Arg>His mutations can confer pH sensitivity to the mutant protein and alter function [28]. We predict that some Arg>His mutations may be adaptive to increased pHi, conferring a gain in pH sensing to the mutant protein.

From our analyses, Arg>His mutations define the mutation landscape of a diverse set of cancers across a range of tissues including brain (low-grade glioma), digestive (colorectal), reproductive (uterine), and blood (AML) cancers. Importantly, these cancers do not have overlapping nucleotide mutation signatures [5], which suggests that the amino acid mutation signatures we identified may reflect other aspects of the cancers including distinct physiological pressures, microenvironment features, or functional requirements. Indeed, these results may help inform studies in the emerging field of Molecular Pathologic Epidemiology (MPE) [29, 30], which seeks to integrate knowledge across disciplines to inform personalized approaches to cancer prevention and therapy. Linking amino acid signatures to physiological or pathological features of the cancer could be important for identifying selective pressures that may be driving or sustaining the cancer as well as for limiting disease progression, particularly where targeted approaches fail [31–33].

## MATERIALS AND METHODS

### Mutation Dataset Filtering

We validated the dataset [5] by comparing known frequencies of well-studied cancer driver genes with observed frequencies in the dataset. Specifically, BRAF is mutated in 40–50% of melanoma samples, and IDH1 is mutated in 75–85%, low-grade glioma, AML, and glioblastoma samples are mutated 75–85%, 8–12%, and 1–5% of the time, respectively. We used the p53 database (http://p53.fr/index.html) to find expected p53 mutation frequency for various cancers: colorectal, head and neck, pancreatic, stomach, liver, and breast cancer have 43%, 42%, 34%, 32%, 31%, and 22% p53 mutation rates, respectively. The observed mutation frequencies were consistently lower than expected for the genes/cancers we assessed, which suggests that the dataset authors [5] were perhaps too stringent in quality control (QC) filtering. Different levels of QC filtering were performed, and we systematically relaxed filters in order to recapitulate the expected mutation frequencies of the selected canonical driver genes. Applying only the ‘sequencing artifact’ QC filter (from [5]) most closely recapitulated expected mutation frequencies for the canonical driver genes, and this filter alone was used for the remainder of the bioinformatics analyses.

### Mapping somatic SNPs

After filtering we used part of the PolyPhen2 [34] pipeline to map mutations to UCSC Canonical transcripts and restricted to nonsynonymous amino acid changes. The following cancers had reduced sample sizes after filtering and nonsynonymous mutation restriction: AML: one sample eliminated through QC filtering, two samples eliminated because all mutations were synonymous; low grade glioma: one sample eliminated because after QC filtering all remaining mutations were synonymous; glioblastoma: two samples eliminated because all mutations were synonymous. All Pilocytic Astrocytoma samples were excluded from future analysis due to low total nonsynonymous mutations per sample.

### Mutation frequency data sets

For the individual sample data, we represent each sample as a row vector with elements giving the mutation counts observed for each nonsynonymous mutation (e.g. Ala>Cys, Ala>Asp, etc.) and removing all samples with <10 total observed mutations. For the aggregated data set, we sum the mutation counts across all samples of the same cancer type (including samples with <10 mutations), giving one row vector for each cancer type where each element represents the total number of observed nonsynonymous mutations across all samples. For non-negative matrix factorization and principal component analysis, we divide each row by the row sum.

NMF is an unsupervised learning method used to decompose a data matrix into a product of two non-negative matrices representing a set of k signals and mixture coefficients. For example if **X** is an *m* × *n* matrix representing the nonsynonymous mutation frequency data, then the NMF of the data is given by

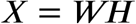

where **W** is an *m* ×*k* matrix with the k columns representing mutation signatures and **H** is a *k* × *n* matrix representing the mixture coefficients that best reconstruct **X**. Often it is not possible to factor **X** exactly, so a typical approach to solving the decomposition will optimize

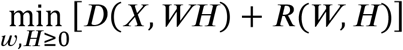

where D() is a loss function (often the Frobenius norm or the Kullback-Leibler divergence) and R() is a regularization function. For our NMF analyses, we utilize the R package *NMF* [35] with default choices for D() and R().

### Principal Component Analysis (PCA)

PCA is a dimension reducing learning method designed to decompose a data matrix into a set of orthogonal bases defined along the major axes of variation within the data. Here we compute the first two principal components from our mutation frequency matrix **X**. The k^th^ principal component is represented by a vector of loadings, *ω*_(*k*)_. The first PC is then calculated as

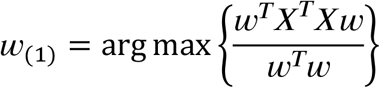

and subsequent PCs are calculated as

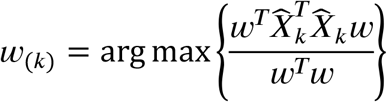

where

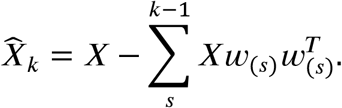

We use the R package *prcomp* to perform all PCA analyses.

### Validation of NMF Mutation Signatures

In order to validate the mutation signatures that we discovered in our data, we sought an orthogonal data set in which to replicate our analysis. We used the COSMIC v81 database of somatic mutations [10]. We first filtered all mutations that were not marked as confirmed somatic mutations. Next, as our original data set (“Alexandrov Data”) had overlapping samples within the COSMIC database, we excluded all samples that were included in our original analysis. Finally, we excluded samples with fewer than 10 total non-synonymous mutations. This filtering resulted in a final data set of 2,236,176 non-synonymous mutations across 15,868 samples. We named this final data set the “COSMIC Data.” We then ran NMF with *k* = 6 signatures on the matrix of individual sample mutation frequencies as described above. Results are shown in S4 Fig. We found that five of the six mutation signatures we originally discovered were replicated in the COSMIC data (Fig 4).

## ACKNOWLEDGEMENTS

This work was supported by National Institutes of Health grants CA178706 to Diane L. Barber and Ryan D. Hernandez; CA197855 to Diane L. Barber, and HG007644 to Ryan D. Hernandez; and National Institutes of Health F32 grant CA177085 to Katharine A. White.

